# Impact of Cesarean Delivery on Reward Behavior and Neurodevelopment in Adult Prairie Vole Offspring

**DOI:** 10.1101/2025.04.01.646614

**Authors:** Katelyn Rogers, Emily Kiernan, Miranda Partie, William Kenkel

## Abstract

Accumulating clinical evidence has shown that birth by Cesarean section (CS) is associated with a higher incidence of disorders involving the dysregulation of dopamine (DA), such as attention deficit-hyperactivity disorder, autism spectrum disorder, and obesity, compared to vaginal delivery (VD). The mesolimbic (ML) system encompasses DAergic neurons that modulate reward processes underlying learning, motivation, and food intake. Previous research has shown that there are lower levels of DA in the prefrontal cortex and higher in the nucleus accumbens (NAc) of CS offspring. Alterations in the ML-DA system as a consequence of birth via CS may impact behavioral response to rewarding stimuli, such as food. Thus, we aimed to ascertain the behavioral and neurodevelopmental outcomes relevant to food reward in CS prairie vole offspring. This study utilized conditioned place preference (CPP) testing to assess learning using context-dependent conditioning, operant conditioning to assess acquisition of a conditioned response and motivation to receive a reinforcer, and immunohistochemistry (IHC) to stain for tyrosine hydroxylase (TH) in the NAc. Behavioral results showed no difference in preference for the conditioned chamber during CPP testing between CS offspring and their VD counterparts. CS prairie vole offspring had a lower average break point during progressive-ratio testing compared to VD offspring, but no difference in response during fixed-ratio 1 or 3 testing. IHC results showed CS offspring had lower levels of TH-immunoreactivity in the NAc core and shell. These findings further support that delivery by CS has long-term neurodevelopmental effects, specifically in the brain’s reward system, and that CS offspring have decreased motivation toward food reward independent of deficits in learning.

## 1. Introduction

Cesarean section (CS) is a surgical procedure that can prevent maternal and infant mortality when medically necessary. The World Health Organization recommends the rate of CS be between 10-15%, stating there is no significant reduction in mortality past this rate; however, as of 2021, the rate of CS is 32.1% of all live births in the United States (World Health Organization, 2018; Osterman et al., 2023). Physician recommendation, maternal request, and policy guidelines contribute to this rate in the absence of indicated medical or obstetric complications under the assumption this mode of birth is without risks (Menacker et al., 2006).

On the contrary, clinical research has shown that being delivered by CS is associated with an increased risk of obesity across development (Li et al., 2013; Darmasseelane et al., 2014; Pei et al., 2014; Yuan et al., 2016). Particularly in childhood, within-family analysis of siblings born either by CS or vaginal delivery (VD) showed CS children have a 64% higher rate of obesity than their VD siblings (Yuan et al., 2016). Additionally, adults born by CS have an at least 33% higher risk of developing obesity (Darmasseelane et al., 2014). We have examined this using an animal model, the prairie vole, and found that CS prairie vole offspring weigh more than their VD counterparts (Kenkel et al., 2024). The impact of obesity at the individual, societal, and economic levels cause concern for public health (Hecker et al., 2022), making it important to understand the biological mechanisms underlying increased risk of obesity in CS offspring.

Animal studies have shown that the mesolimbic dopaminergic (ML-DA) system, the neural system implicated in reward-mediated behavior, is altered in CS rodent offspring (El-Khodor & Boksa, 1997; Wise, 2005). After birth, there are lower levels of DA and DA metabolites in the brain of CS pups suggesting being delivered by CS alters neurotransmitter exposure around the sensitive period of birth (Ikeda et al., 2019). In adult rats delivered by CS, there are lower levels of DA in the prefrontal cortex (PFC) and higher in the nucleus accumbens (NAc) compared to VD offspring, suggesting delivery by CS has long-term neurodevelopmental effects on the ML-

DA system (El-Khodor & Boksa, 1997). Clinical research has shown similar alterations in the ML-DA system in obese patients impacted by reward sensitivity to food (Volkow et al., 2011). Preclinical research has shown that adult obesity prone rats have higher levels of extracellular DA in the NAc associated with increased motivation to receive a high-fat pellet during an operant task (Narayanaswami et al., 2013). Alterations to the ML-DA system as a consequence of being born by CS has yet to be explored as a mechanism for the associated increased risk of obesity despite the implications of DA and body weight on behavioral response to rewarding stimuli, such as food.

In this study, we aimed to ascertain the behavioral and neurodevelopmental outcomes relevant to food reward in adult CS prairie vole offspring. Conditioned Place Preference (CPP) testing was used to assess learning using context-dependent conditioning, operant conditioning was used to assess acquisition of a conditioned response and motivation to receive a reinforcer, and immunohistochemistry was used to stain for tyrosine hydroxylase-immunoreactivity (TH-ir) in the NAc between CS and VD offspring. We anticipated CS offspring would 1) spend more time in the conditioned chamber during CPP testing, 2) have a higher rate of response in all schedules during operant conditioning, and 3) have higher levels of TH-ir in the NAc compared to VD offspring. The results from this study further support that delivery via CS has long-term neurodevelopmental effects which may contribute to decreased motivation toward food reward seen during operant testing.

## 2. Methods

This study utilized laboratory-bred prairie voles (*Microtus ochrogaster*). All subjects were given ad libitum access to food and housed at room temperature (23°C) on a 14:10 light:dark cycle. On post-natal day (PND) 21, pups were weaned and housed in groups of 2-4 with same-sex, same-condition litter mates. Each offspring condition received a mixed high-fat diet of standard prairie vole chow (LabDiet Rabbit Diet HF 5326) and standard mouse chow (LabDiet Prolab RMH 3000). Subjects underwent CPP testing on PND 41-45 and operant testing on PND 60-76, brain tissue was collected on PND 77 (Fig. 1A). All procedures in this study were approved by the University of Delaware Institutional Laboratory Animal Care and Use Committee and were conducted in accordance with the National Institutes of Health Guide for the Care and Use of Laboratory Animals.

**Fig. 1.**
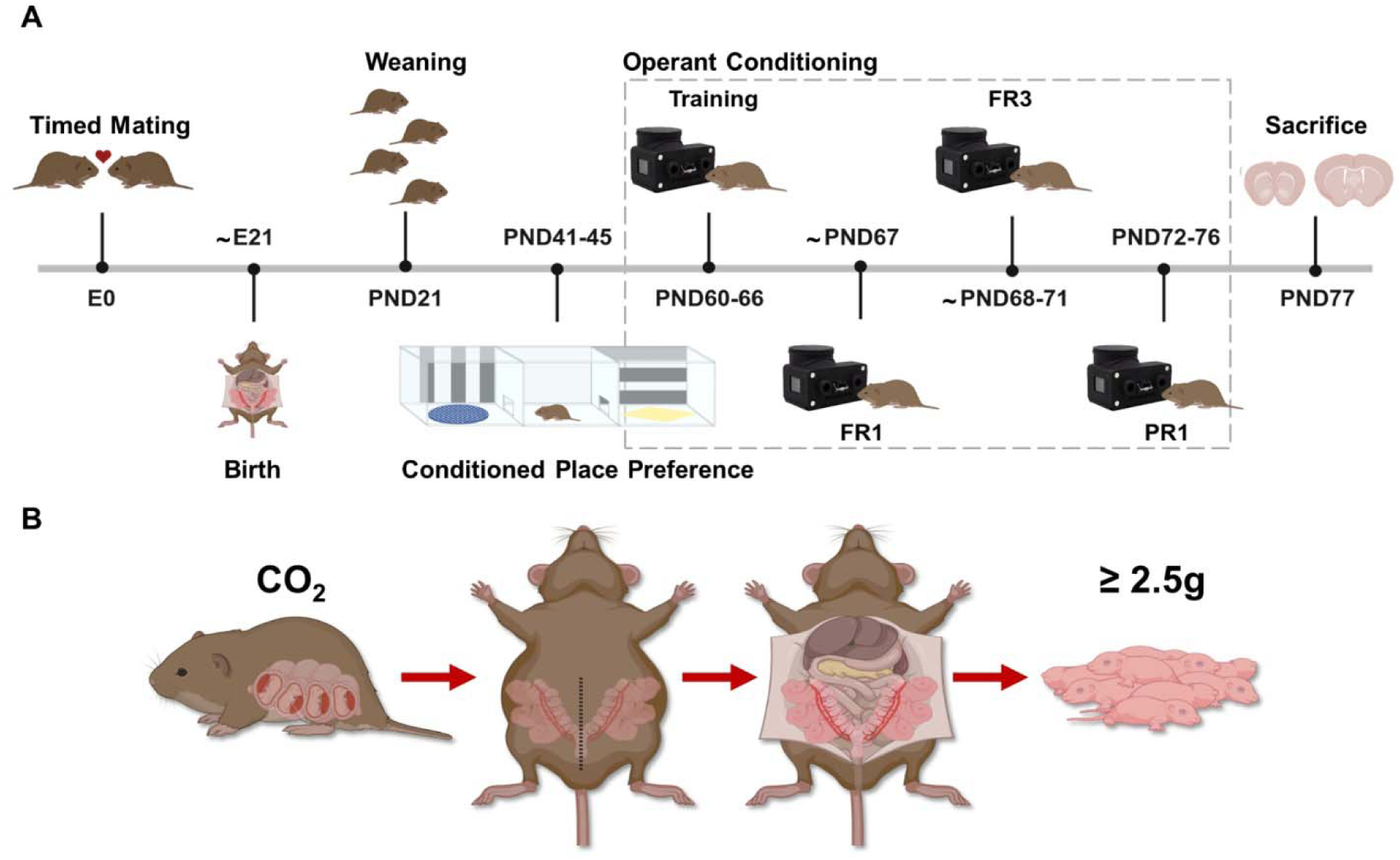
A) Experimental timeline of all procedures conducted in this study. Animals were generated based on a timed mating protocol and then dams either delivered via Cesarean section (CS) or vaginal delivery (VD) around embryonic day 21. On post-natal day (PND) 21, subjects were weaned and housed with same-sex, same-condition littermates. Conditioned Place Preference testing occurred on PND 41-45. Operant conditioning training and testing occurred on PND 60-76. Subjects underwent 6 days of home cage training and then 1-3 days of fixed-ratio 1 (FR1) testing. Subjects that met the criterion (≥20 nose pokes) were run through 2-4 days of fixed-ratio 3 (FR3) testing and 5 days of progressive-ratio 1 (PR1) testing. Sacrifice occurred on PND 77, and brain tissue was extracted for histology. B) CS procedure. Dam was exposed to CO_2_ for 90 seconds then cervically dislocated. Pups were removed via laparotomy. Created in BioRender. Rogers, K. (2025) https://BioRender.com/w35w174

### 2.1. Birth Mode

Subjects were generated based on a previously established timed mating protocol (Kenkel et al., 2023; Kenkel et al., 2024) to reduce the chance of prematurity and increase the chance of predicting time of delivery. CS deliveries occurred around the time of expected delivery, embryonic day ∼21.5. In the CS condition, the dam was exposed to CO_2_ for 90 seconds to avoid the neurodevelopmental consequences of pharmacological anesthesia such as apoptosis of developing neurons and hypoxia in the fetal brain (Anand & Soriano, 2004; Castillo-Ruiz et al., 2018) and then cervically dislocated. Pups were removed via laparotomy (Fig. 1B). In the VD condition, pups were born by spontaneous delivery. It is important to note that the CS procedure was non-survival and that CS litters were given to foster parents because female voles who deliver by CS are non-maternal (Hayes & De Vries, 2007). We controlled for the potential effects of differences in rearing of non-biological pups (Perkeybile et al., 2013; Perkeybile et al., 2015) by also cross fostering VD litters. Litters were culled to 3-5 pups and had an average weight ≥2.5 grams per pup before being placed with foster parents.

### 2.2. Weights

Offspring from both groups were weighed every 7 days starting on PND 21 until sacrifice on PND 77.

### 2.3. Conditioned Place Preference

CPP testing (PND 41-45) consisted of a three-chamber apparatus (∼50cm x 100cm) with a neutral center chamber and openings on either side to allow access to two contextually different chambers. The left and right compartments contained distinct textured silicon mats (raised bumps or holes) and wallpaper (vertical or horizontal black/white stripes). Orientation of mats and wallpaper were counterbalanced between subjects. On day one of testing, subjects were placed in the center chamber and allowed to freely explore all chambers for 10 minutes to establish baseline preference for either chamber. Videos were hand scored for time spent in each chamber, excluding the middle chamber, to determine which was preferred and nonpreferred. On days 2-4 of testing, we conditioned against the preference by confining subjects in the nonpreferred chamber (conditioned chamber) with 3 mm sprinkles (Sugar Pearls, CakeDeco) in a petri dish for 15 minutes, and then 3-4 hours later confined the subject to the preferred compartment paired with nothing for 15 minutes. A plastic board was used to confine subjects to a chamber and the order of conditioning was counterbalanced across days 2-4. On day 5 of testing, subjects were placed in the center compartment of the chamber and allowed to freely explore for 10 minutes. Videos recorded on day 1 and 5 of CPP testing were uploaded into idTracker, an automated video tracking software (Pérez-Escudero et al., 2014), to identify individual subjects and measure time spent in each chamber. Preference score for the conditioned chamber was determined by subtracting time spent in the nonpreferred chamber during baseline testing on day 1 from time spent in the conditioned chamber during testing on day 5.

### 2.4 Operant conditioning

Training of the conditioned response (nose poke) occurred on PND 60-66 in the home cage. The Feeding Experimentation Device 3 (FED3), an open-source automated pellet dispensing system (Matikainen-Ankney et al., 2020), was placed inside the home cage on day 1 for 30 minutes using an automated method of magazine training to establish an association between the device and food. This method used the free feeding FED3 paradigm where it automatically dispensed a grain pellet (TestDiet MLab Rodent Tablet 1811143, 20mg) when it sensed the removal of the previous pellet from the pellet well. After magazine training, the FED3 remained in the home cage for the remainder of training on a fixed-ratio 1 (FR1) schedule where it dispensed one grain pellet at the onset of one nose poke in the target sensor. A tone and blue light were paired with the response.

On PND 67, subjects were removed from the home cage and placed individually into a small polycarbonate cage with wood chip bedding (Beta Chip Hardwood) and ad libitum access to only water for 5 hours. This food deprivation procedure was followed for all schedules conducted during testing and designed to reduce significant loss in body weight while increasing the likelihood that subjects would perform the operant during testing. The FED3 was placed in the cage on a FR1 schedule for 30 minutes. Subjects were required to provide ≥20 nose pokes in the target sensor during testing to demonstrate acquisition of the conditioned response and mastery of the device. If subjects failed to meet criterion during the first FR1 testing period, they were run up to an additional two days on a FR1 schedule to meet criterion before being removed from testing. One subject per sex per litter that met the criterion were run through FR3 (2-4 days) and progressive ratio 1 (PR1) (5 days) schedules for 30 minutes each day. FR3 required three responses to receive one grain pellet. The transition in required responses between FR1 and FR3 testing was used to assess acquisition measured by the rate of response provided during testing. PR1 required an exponentially increasing number of responses by 1 (1, 2, 3, etc.) to receive one grain pellet to assess motivation as a function of breakpoint, the maximum rate of response that a subject would give to receive an absent reinforcer. Termination of the testing period required the subject to stop responding within 10 minutes of the last response.

Subject’s responses in both sensors were logged by the FED3. Responses recorded in the target sensor for FR1 testing were analyzed for all subjects that met the criterion. Responses recorded in the target sensor during FR3 testing were averaged across the last two test days. Breakpoint was calculated from responses recorded in the target sensor during PR1 testing, then averaged across the last three test days.

### 2.5. Tissue Extraction and Immunohistochemistry

Sacrifice occurred on PND 77. Subjects were exposed to isoflurane for ∼5 minutes and then rapidly decapitated. Brains were extracted from subjects and placed free-floating in 4% paraformaldehyde for 24 hours then switched to a 25% sucrose and 0.01% sodium azide solution for storage at 4°C. Fixed tissue was sectioned using a cryostat microtome at 30 µm from medial PFC to ventral hippocampus and stored in cryoprotectant at -20°C.

Tissue was rinsed in 0.05M potassium phosphate buffered saline (KPBS) for 5 minutes three consecutive times to remove cryoprotectant, then incubated in 1% hydrogen peroxide for 30 minutes. Next, the tissue was rinsed in KPBS for 5 minutes three consecutive times. After this, tissue was incubated in primary mouse TH antibody (MilliporeSigma antibodies # IHCR10056; 1:1000 dilution in KPBS) for 60 minutes at room temperature and then incubated for 48 hours at 4°C. Sections were then rinsed for 5 minutes three consecutive times in KPBS and then incubated in 1% horse serum (Thermo Fisher Scientific) + KPBS + 0.4% Triton-X (KPBS_TX_) for 30 minutes at room temperature. The tissue was then incubated in horse anti-mouse/rabbit IgG secondary antibody (Vector Labs; 1:600 dilution in KPBS_TX_) for 60 minutes at room temperature then rinsed for 5 minutes five consecutive times in KPBS. Next, tissue was incubated in avidin-biotin peroxidase complex (Vector Labs; Vectastain Elite ABC Universal kit; 45µl A, 45µl B per 10 ml KPBS_TX_) for 60 minutes at room temperature and then rinsed in KPBS for 5 minutes three consecutive times. The tissue was then rinsed in 0.175M sodium acetate for 5 minutes three consecutive times and then placed in a nickel-diaminobenzene solution (1 tablet diaminobenzene, 50 ml sodium acetate, 1.25g nickel sulfate) per dish. Hydrogen peroxide (41.5 µl per 50 ml) was added immediately before incubation in the nickel-diaminobenzene solution for 8 minutes at room temperature. Finally, tissue was rinsed in sodium acetate for 5 minutes three consecutive times and then KPBS for 5 minutes three consecutive times.

Brain sections were mounted onto superfrost plus microscope slides (Thermo Fisher Scientific), dehydrated in ascending ethanol dilutions, cleared with Histoclear, and then cover- slipped. Slides were then scanned using the EVOS M7000 microscope using a 4x objective. Tiled images were stitched together to form one image of the slide. Individual brain sections were warped to fit the Vole Brain Atlas overlays 19-21 (Yee et al., 2016) using the BigWarp plugin in ImageJ (Schneider et al., 2012) to outline the NAc core, NAc shell, and Caudate-Putamen (CP) for all subjects. Warped images were then used to measure TH-ir by pixel intensity per target region using custom MATLAB code. Pixel intensity of the NAc core, NAc shell, and CP were collapsed across atlas pages (anterior to posterior).

### 2.6. Data Analysis

All statistical analyses were performed using RStudio. Outlying data was identified by calculating the lower and upper limit for outliers, then removed. Assumptions of normality and homogeneity of variance were tested using Shapiro-Wilk’s test and Levene’s test, respectively. Repeated measures linear mixed effects modeling was used to make comparisons between CS and VD offspring (adult body weights), repeated over individual subjects or litter, using the lmer() function from the *lme4* library (Bates et al., 2015) followed by performing an anova on the model (type III sum of squares). Linear mixed effects modeling was used for non-repeated measures (FR3, PR1, IHC). Birth mode and sex were considered as main effects. In the case of significant main or interaction effects, post-hoc analyses were performed by either varying the levels of the grouping factor and/or making comparisons for each individual time point for repeated measures. Data was analyzed by Kruskal-Wallis and Mann-Whitney U tests when data was not normally distributed (litter weight, CPP, FR1). Sample sizes are listed for each measure and data are reported as either means plus/minus standard error or medians, alpha = 0.05, and effect sizes are reported as Cohen’s *d*.

## 3. Results

### 3.1. Weights

The median litter weight of CS offspring (n = 13) was 13g (IQR = 2.52) and the median litter weight of VD offspring (n = 17) was 13.3g (IQR = 2.82) at time of delivery/discovery. There was no significant difference in litter weight as a function of birth mode (p = 0.38; Fig. 2A).

**Fig. 2.**
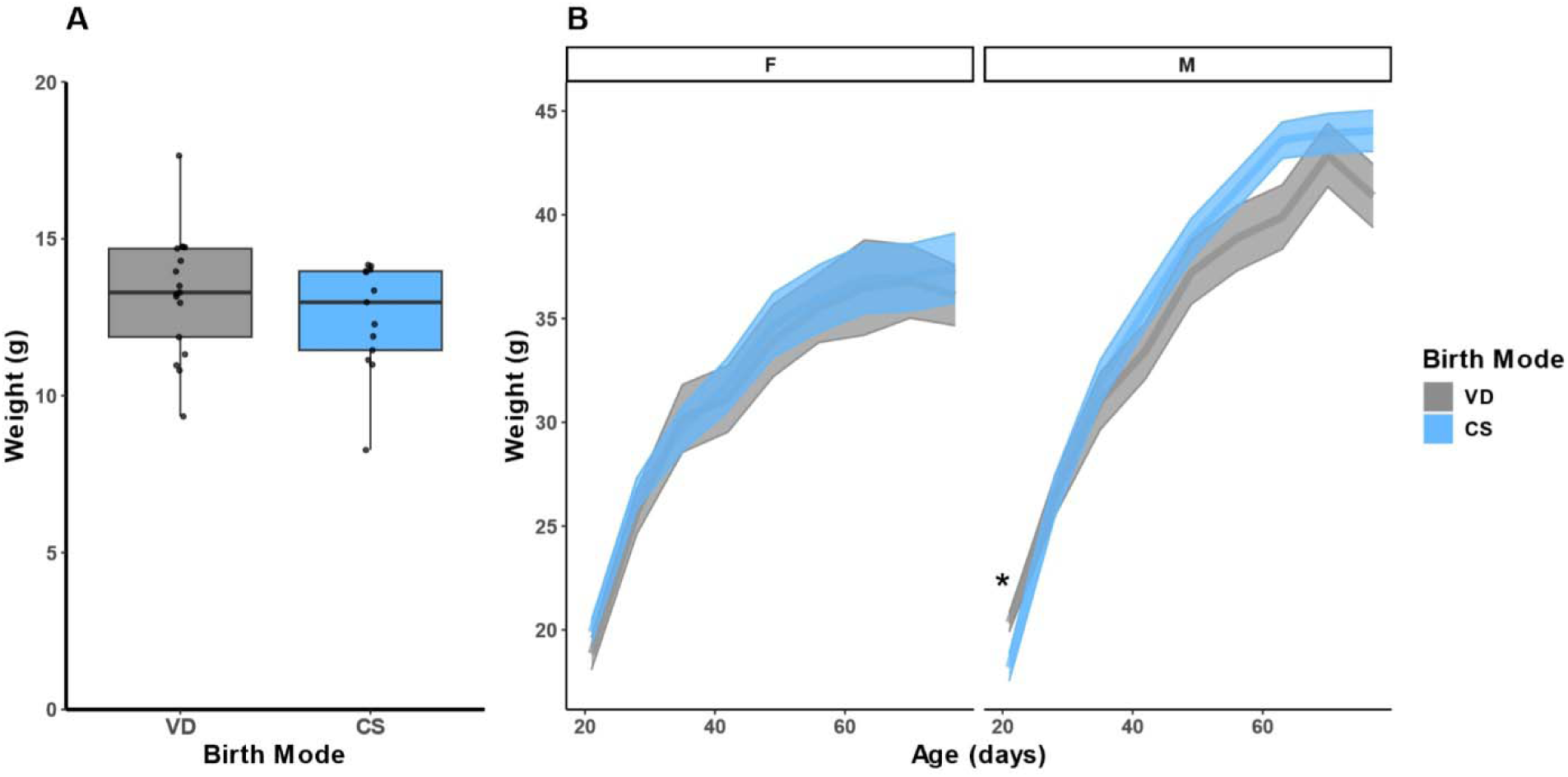
A) Median litter weight (g), measured immediately after birth or at time of delivery, between prairie vole pups delivered by Cesarean section (CS) or vaginal delivery (VD). There was no significant difference in litter weight as a function of birth mode (p = 0.38). B) Average weight (g) ± standard error between prairie vole offspring born by CS or VD from post-natal day (PND) 21 to 77. Weights were taken every seven days starting on PND 21 until sacrifice from a subset of subjects run through operant testing. There was a significant main effect of age (p < 0.001) and sex (p = 0.19), and an interaction between age and birth mode (p = 0.029) and age and sex (p < 0.001). There was no difference in weight as a function of birth mode (p = 0.44). Post-hoc analyses revealed there was an interaction between birth mode and sex at PND 21 where VD males weighed more than CS males (p = 0.017; * p < 0.05).

Adult body weight measures included a subset of subjects run through operant testing (CS n = 34; VD n = 24). One outlying data point was removed from the CS group on PND 28. Results showed a significant effect of age (F(1, 446.13) = 1592.136, p < 0.001) and sex (F(1, 135.75) = 5.63, p = 0.019), but no significant effect of birth mode on body weight (p = 0.44).

There was an interaction of age and birth mode (F(1, 446.13) = 4.78, p = 0.029) and a significant interaction of age and sex (F(1, 446.18) = 47.84, p < 0.001) with VD and males contributing to increasing weight across time. Post-hoc analysis showed there was an interaction between mode and sex (F (1, 54) = 6.045, p = 0.017) at PND 21 where VD males weighed more than all other groups. There was a main effect of sex at PND 42 (F(1,54) = 5.83, p = 0.019, d = -0.65), PND 49 (F(1, 54) = 6.85, p = 0.011, d = -0.705), PND 56 (F(1, 54) = 9.52, p = 0.003, d = -0.83), PND 63 (F(1, 46) = 13.47, p < 0.001, d = -1.01), PND 70 (F(1,49) = 19.021, p < 0.001, d = -1.23), and PND 77 (F(1, 54) = 16.63, p < 0.001, d = -1.082) on body weight where males weighed more than females (Fig. 2B).

### 3.2. Conditioned Place Preference

21 CS and 24 VD offspring were run through CPP testing. The median preference score of CS offspring was 89.3 (IQR = 324) and the median preference score of VD offspring was 37.8 (IQR = 197). There was no significant effect of sex or birth mode on preference score (p = 0.9; Fig. 3). Both CS and VD subjects spent more time in the conditioned chamber on test day 5 (p = 0.027; data not shown) compared to baseline testing, determining both groups formed a significant preference for the conditioned chamber.

**Fig. 3.**
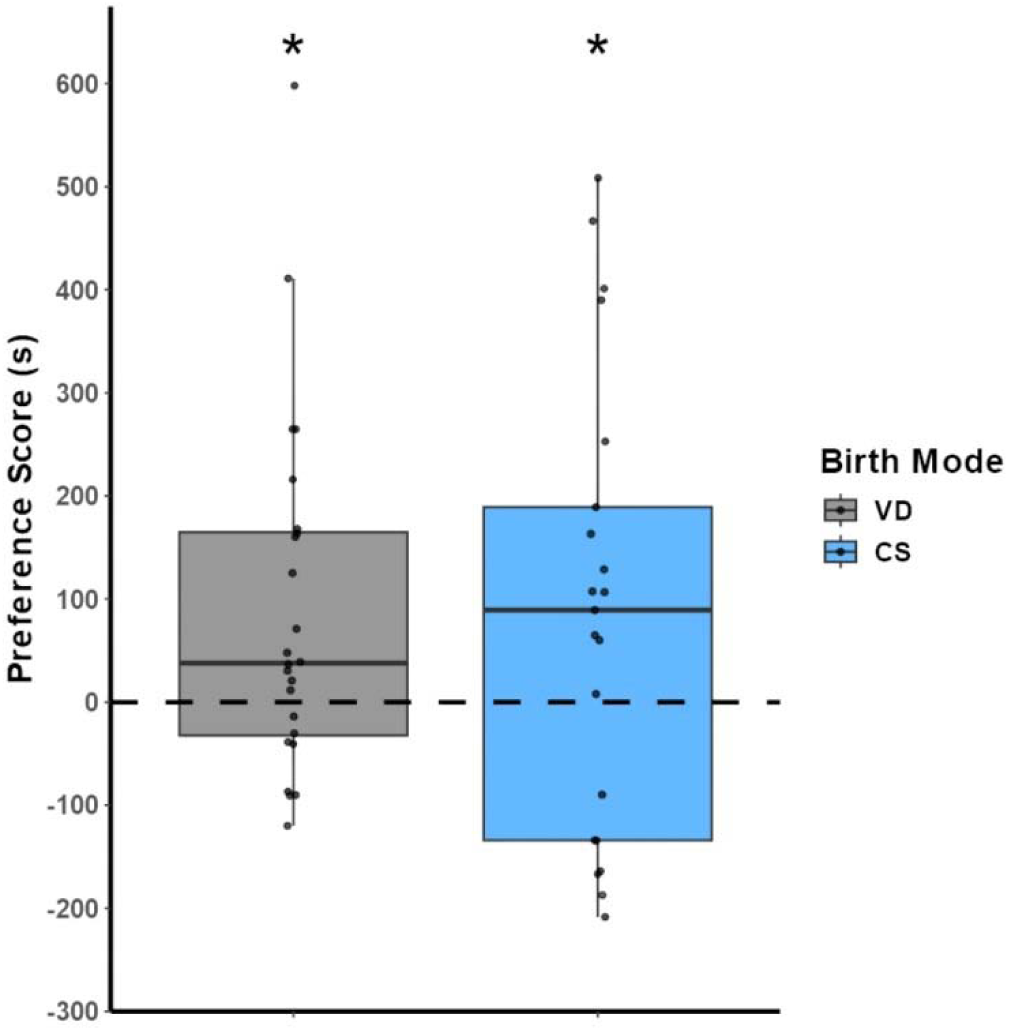
Median preference score (s) for the conditioned chamber during Conditioned Place Preference testing between prairie vole offspring delivered via Cesarean section (CS) or vaginal delivery (VD). Preference score was calculated as the difference in time between day 5 testing (post-conditioning) and baseline testing (pre-conditioning). There was no significant effect of birth mode on preference score (p = 0.9). Both CS and VD subjects spent more time in the conditioned chamber on test day 5 (p = 0.027; * p < 0.05).

### 3.3. Operant Conditioning

37 subjects (CS n = 16, VD n = 21) out of 93 total subjects (CS n = 35, VD n = 58) run through FR1 testing met the criterion (∼40% total success rate). Of those subjects that met the criterion, the median response of CS offspring was 35 (IQR = 17.8) and the median response of VD offspring was 29 (IQR = 15) during FR1 testing. There were no effects of birth mode or sex on response (p = 0.69, Fig. 4A). 13 CS and 18 VD offspring were included in FR3 testing because one outlying data point was removed from both the VD and CS male groups. There was no effect of day on response (p = 0.61), therefore researchers collapsed across the last two test days. There were no effects of birth mode or sex on response during FR3 testing (p = 0.26, Fig. 4B). For PR1 testing, 14 CS and 19 VD offspring were included. There was no effect of day on response (p = 0.72), therefore researchers collapsed across the last three test days. There was no effect of sex on break point (p = 0.59); however, CS prairie vole offspring had a lower average breakpoint compared to their VD counterparts (F(1, 29) = 6.25, p = 0.018, d = -0.45; Fig. 4C). Due to issues with the FED3 during testing, two subjects did not have their response logged during FR1 testing, four subjects did not have their response logged during FR3 testing, and one subject did not have their response logged during PR1 testing.

**Fig. 4.**
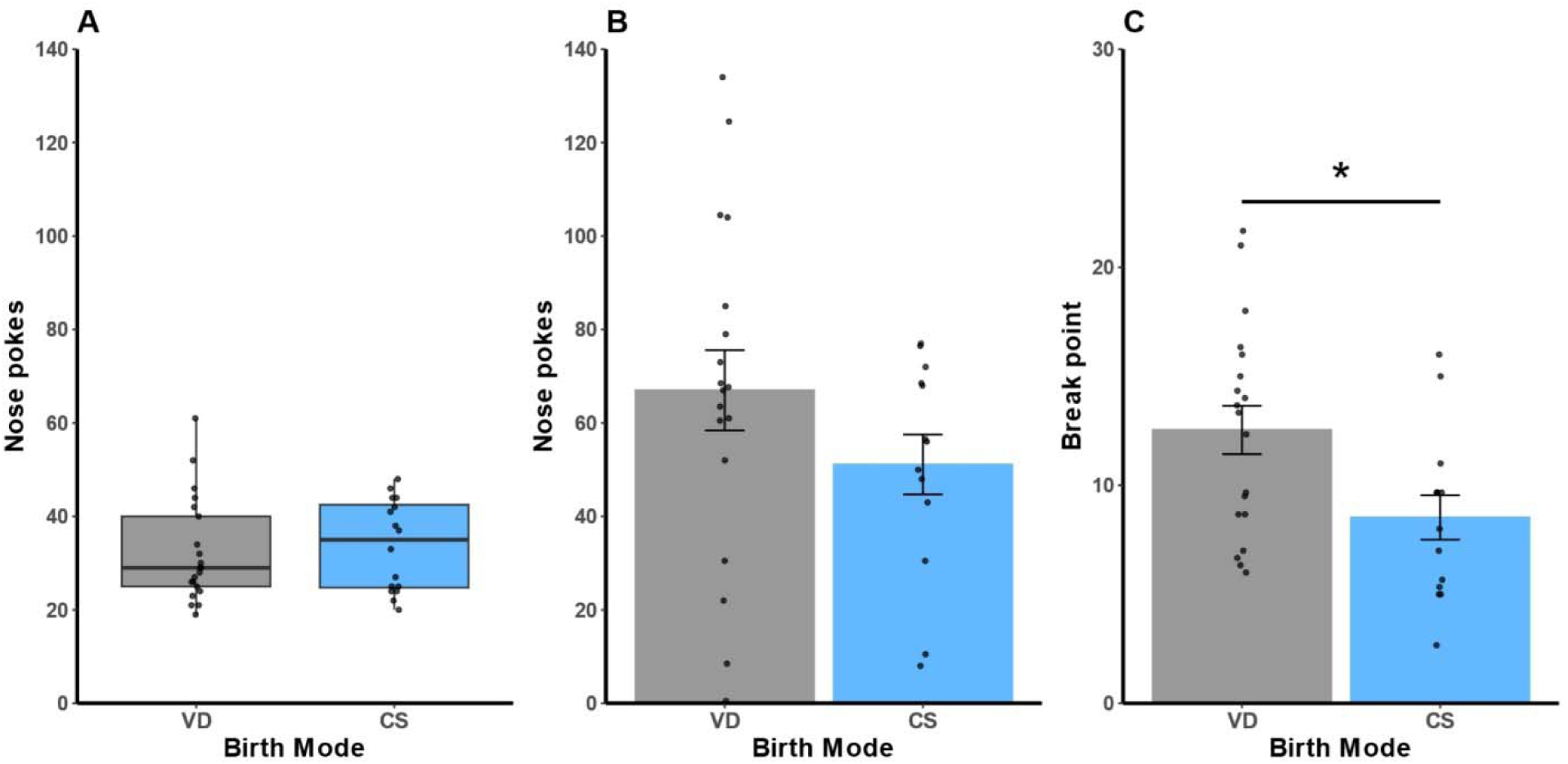
A) Median nose pokes during fixed-ratio 1 (FR1) testing between prairie vole offspring delivered via Cesarean section (CS) or vaginal delivery (VD). There was no effect of birth mode on response during FR1 testing (p = 0.69). B) Average nose pokes ± standard error during fixed-ratio 3 (FR3) testing between CS and VD prairie vole offspring. There was no effect of birth mode on response during FR3 testing (p = 0.26). C) Average break point ± standard error during progressive-ratio 1 (PR1) testing between CS and VD offspring. CS prairie vole offspring had a lower average break point during PR1 testing compared to VD offspring (p = 0.018; * p < 0.05).

### 3.4. Immunohistochemistry

11 VD subjects and 10 CS subjects were included in the TH-ir analysis. Researchers collapsed across position (anterior-posterior) for each target region since there was no effect of position on TH-ir (NAc core p = 0.47; NAc shell p = 0.92; CP p = 0.3). Results showed a main effect of birth mode where CS prairie vole offspring had lower levels of TH-ir in the NAc core (F(1, 17) = 4.53, p = 0.048, d = 0.93; Fig. 5C) and shell (F(1,16) = 7.05, p = 0.017, d = 1.207; Fig. 5D), but no significant effect in the CP (p = 0.25; Fig. 5E). There was no significant main effect of sex on TH-ir for the NAc core (p = 0.34), NAc shell (p = 0.62), or CP (p = 0.054).

**Fig. 5.**
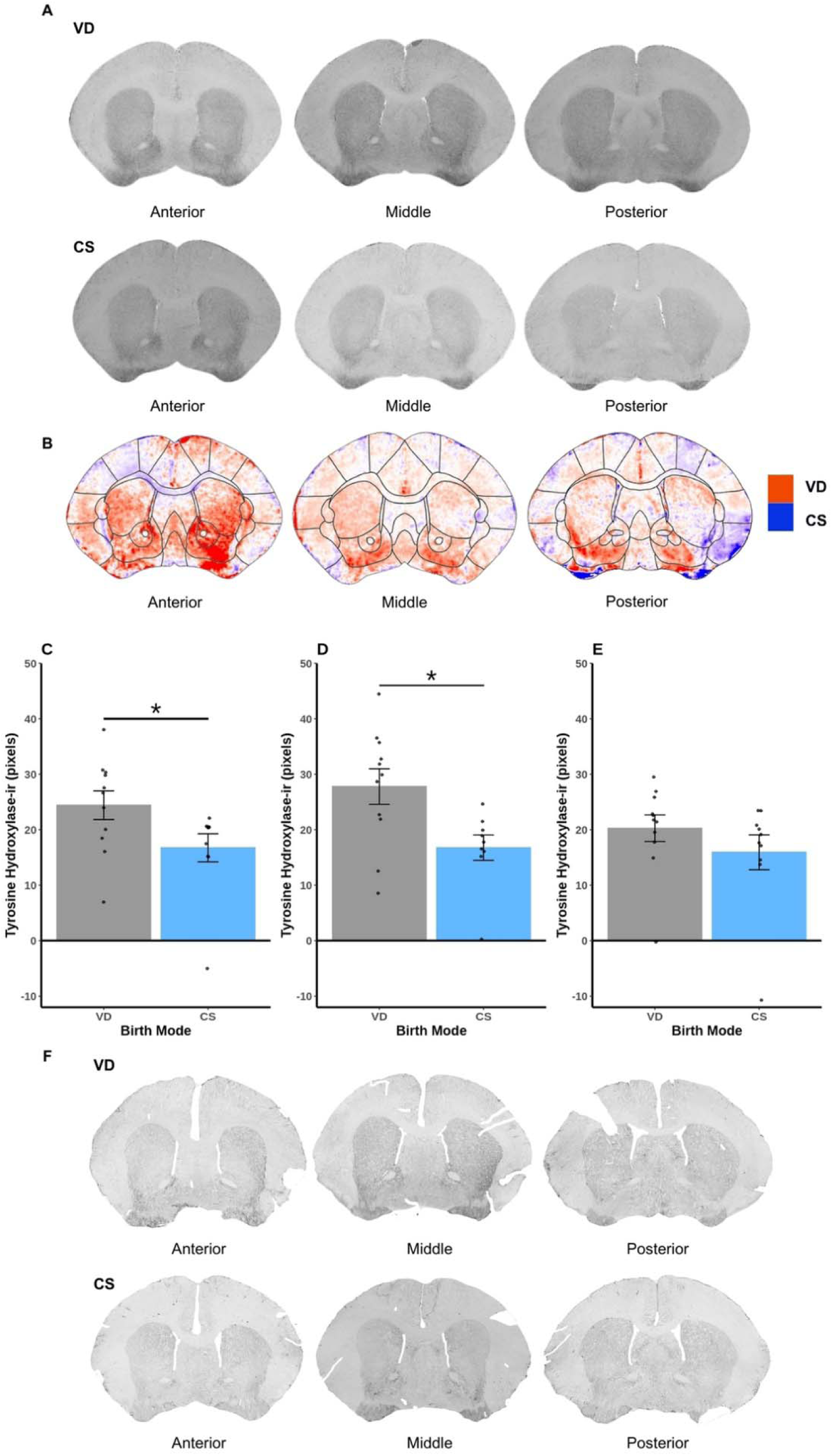
A) Composites of warped brain section images between prairie vole offspring delivered via Cesarean section (CS) and vaginal delivery (VD) (anterior to posterior). B) Heat maps of pixel intensity of tyrosine hydroxylase-immunoreactivity (TH-ir) (anterior to posterior) between CS (blue) and VD (red) offspring. C) Average TH-ir (pixels) ± standard error in the nucleus accumbens (NAc) core (collapsed anterior to posterior) between CS and VD offspring. CS prairie vole offspring had lower levels of TH-ir in the NAc core compared to VD offspring (p = 0.048; * p < 0.05). D) Average TH-ir (pixels) ± standard error in the NAc shell (collapsed anterior to posterior) between CS and VD offspring. CS prairie vole offspring had lower levels of TH-ir in the NAc shell compared to VD offspring (p = 0.017; * p < 0.05). E) Average TH-ir (pixels) ± standard error in the Caudate-Putamen (CP) (collapsed anterior to posterior) between CS and VD offspring. There was no effect of birth mode on TH-ir in the CP (p = 0.25). F) Anterior to posterior representative photomicrographs of warped brain sections stained for TH split by birth mode.

## 4. Discussion

The main findings in this study are that adult CS prairie vole offspring have decreased motivation toward food reward and lower levels of TH-ir in the NAc core and shell compared to VD offspring. We were able to demonstrate that CS birth has implications for long-term neurodevelopment and behavioral response toward food, independent of weight or acquisition of the conditioned response, suggesting birth mode impacts reward value of appetitive stimuli.

Previous work from our group showed that CS prairie vole offspring weigh more than VD controls in adolescence and adulthood (Kenkel et al., 2024). The use of prairie voles as our model organism comes from accumulating evidence supporting that voles are better suited to endure conventional laboratory housing conditions compared to traditional rodent models of obesity, such as mice. Vole thermoneutral conditions (∼25-30°C) are closer to ambient temperature (23°C) compared to mice (30-32°C) (Beck & Anthony, 1971; Hylander & Repasky, 2016). Previous work from our group demonstrated that prairie voles show minimal change in thermoregulatory function, such as heart rate (∼6%), between 18°C and 30°C, whereas mice show a twofold increase in heart rate outside of thermoneutrality (300 to 600 beats/min from 30°C to 20°C) (Kenkel et al., 2024; Swoap et al., 2008). The increase in heart rate illustrates the metabolic load required to maintain thermoneutrality under conditions of cold stress (Maloney et al., 2014; Swoap et al., 2004; Swoap et al., 2008), even making it more difficult for mouse models of obesity to gain weight in traditional housing temperatures (Feldmann et al., 2009; Stemmer et al., 2015). In this study, we were unable to replicate previous findings demonstrating CS offspring weigh more than VD offspring in adulthood (Kenkel et al., 2024).

This could be due to sampling from a subset of subjects run through operant conditioning resulting in smaller sample sizes. Any difference in weight gain between groups would likely not result from weight at time of birth/discovery as we found no difference in litter weight (Fig. 2A).

In this study, we used CPP testing to assess the association of contextual environment with an appetitive stimulus measured by difference in time spent in a positively reinforced chamber before and after conditioning. Results show that there was no difference in preference for the conditioned chamber between CS and VD offspring, suggesting there was no effect of birth mode on the reward value of sucrose that would enhance or diminish the context-dependent association. Regardless, both groups were able to develop a CPP for sucrose (Fig. 3). It has previously been shown that lesioning the NAc in rats diminishes their ability to form context-dependent associations (Ito et al., 2008), thus based on previous literature demonstrating higher levels of DA in the NAc we anticipated CS offspring to have a higher preference for the conditioned chamber during CPP testing. Further investigation of the reward value of sucrose between CS and VD subjects needs to be done. Prior work from our group has shown there is no difference in preference between CS and VD offspring using traditional two-bottle home cage sucrose preference testing (Kenkel et al., 2024). We intend on extending this by using sucrose preference testing with an automated two-bottle lickometer, the Sipper (Godynyuk et al., 2019), to get more precise individual measures of preference between groups compared to traditional home cage measures.

We used operant conditioning to assess acquisition of the conditioned response (nosepokes) and motivation for the reinforcer (grain pellets) measured by responses during FR1 and FR3 testing and break point during PR1 testing. There was no difference in the number of responses between groups during FR1 or FR3 testing (Fig. 4A, Fig. 4B); however, CS adult prairie vole offspring had a lower average breakpoint during PR1 testing compared to VD controls (Fig. 4C). This suggests that adult CS offspring have decreased motivation towards food reward, contrary to our prediction. Results from FR1 and FR3 testing suggest that the break point findings are not a result of difficulty acquiring the conditioned response. Additionally, higher body mass in adult rats is associated with increased motivation to receive a high-fat pellet during a PR1 operant task (Narayanaswami et al., 2013). Results from this cohort showed no significant differences in body weight between groups (Fig. 2B), suggesting that PR1 results are not due to this association, but from being delivered via CS. The PR1 results are consistent with prior observations that adult CS prairie vole offspring consume less chow than VD offspring, measured as collective home cage food consumption over a 24 hour period at PND 70 (Kenkel et al., 2024). This study benefited from individual measures of response for food reward during multiple days of FR1, FR3, and PR1 testing compared to our previously published food consumption measures (Kenkel et al., 2024). This is the first study of operant conditioning in voles using a novel operant conditioning protocol and the FED3. With that in mind, one caveat to this measure is the success rate (∼40%) of subjects who were able to meet the criterion (≥20 nose pokes) during FR1 testing. We intend on improving our current methods by moving away from home cage training and toward individual training with manual shaping to aid in the success of training subjects and further assessing acquisition of the conditioned response. Second, we intend on extending FR1 training to account for differences in acquisition, as our protocol included 6 days of home cage training whereas other studies of operant conditioning using prairie voles include up to 20 days of training with food reward (Beery et al., 2021; Vahaba et al., 2022). Future studies would benefit from the inclusion of a fixed-ratio 5 schedule, which requires five responses to receive one reinforcer, after FR3 testing to further assess motivation and learning after CS delivery in adult prairie voles. Lastly, future studies would benefit from evaluating the behavioral response toward the reinforcer during operant testing by including latency to nose poke and inter-response rate intervals as additional measures of acquisition, motivation, and feeding behavior.

Prior work has shown that adult CS rat offspring have higher levels of DA in the NAc as measured by high-pressure liquid chromatography (El-Khodor & Boksa, 1997). We sought to investigate further based on these findings and used IHC to stain for TH, the rate limiting enzyme of DA synthesis, to assess TH-ir in the NAc and CP. Contrary to our prediction, we found that adult CS prairie vole offspring had lower levels of TH-ir compared to VD offspring in the NAc core and shell (Fig. 5C, Fig. 5D). Differences in methodology and/or animal model used to assess TH could explain this discrepancy in results. There was no significant difference in TH-ir in the CP between groups, which suggests that the implications of CS birth on TH were specific to the reward system, particularly the NAc. We were able to further support that being delivered by CS has long-term neurodevelopmental effects on TH in the NAc. Considering the role of DA in motivated behaviors (Olds & Milner, 1954; Wise, 2005; Wise, 2006), our IHC results may explain the decreased motivation toward food reward observed in CS offspring during PR1 testing. This dysregulation in TH could arise from alterations in developmental programming in early life as a result of changes in hormone or microbe exposure at or around the time of birth, which may be important for typical brain DA development and behaviors. CS rat pups have lower levels of plasma epinephrine measured immediately after birth (El-Khodor & Boksa, 1997). Adult CS rat offspring have higher levels of TH in the NAc after repeated mild stress in adulthood and an enhanced response to amphetamine -induced locomotion (El-Khodor & Boksa, 1998; Boksa & Zhang, 2008). Exogenous epinephrine subcutaneously injected immediately after birth ameliorates amphetamine -induced hyperactivity and normalizes altered TH activity in the NAc of adult CS born rats compared to CS offspring without acute epinephrine injections and VD controls (Boksa & Zhang, 2008). These findings suggest alterations in exposure to epinephrine around the time of birth contribute to long-term regulation of DA. While results from the present study further implicate CS in dysregulation of TH persisting into adulthood, one limitation of this measure is its small sample sizes. Future work would benefit from larger sample sizes. Additionally, more work needs to be done investigating the role of hormones at the time of birth on development of the brain’s reward system and subsequent reward-mediated behavioral outcomes in CS offspring.

In conclusion, results from this study showed decreased motivation toward food reward and lower levels of TH-ir in the NAc in adult CS offspring compared to VD counterparts demonstrating that birth mode can impact long-term neurodevelopment and subsequent reward-mediated behavior. Considering our findings and epidemiological evidence associating birth by CS with disorders involving the dysregulation of DA, including attention deficit-hyperactivity disorder and autism spectrum disorder, we contend that the broader neural and behavioral implications of DA dysregulation as a consequence of CS warrants further investigation. It will be of interest to determine factors around birth, including exposure to hormones and/or the microbiome, which may contribute to altered development in early life leading to dysregulation of DA in adulthood.

## Acknowledgements

The authors would like to thank the contributions of the University of Delaware Office of Laboratory Animal Management, Nick Foley for his help building the FED3s, and Dr. Gwen Talham in particular. This work was supported by funding from the following sources: R01HD111737 (WMK), P20GM103653 (WMK)

## Declarations of interest

**none**

## Notes

### Competing Interest Statement

The authors have declared no competing interest.

